# Impairments in hippocampal oscillations accompany the loss of LTP induced by GIRK activity blockade

**DOI:** 10.1101/2023.05.05.539539

**Authors:** Ana Contreras, Souhail Djebari, Sara Temprano-Carazo, Alejandro Múnera, Agnès Gruart, José M. Delgado-Garcia, Lydia Jiménez-Díaz, Juan D. Navarro-López

**Author notes:** **Correspondence to:** J.D. Navarro-López and L. Jiménez-Díaz, NeuroPhysiology and Behavior Lab, Universidad Castilla-La Mancha, 13071-Ciudad Real (Spain). These authors share first authorship (Ana Contreras, Souhail Djebari). These authors contributed equally to this work and share last authorship (Lydia Jiménez-Díaz, Juan D. Navarro-López).

## Abstract

Learning and memory occurrence requires of hippocampal long-term synaptic plasticity and a precise neural activity orchestrated by brain network oscillations, both processes reciprocally influencing each other. As G-protein-gated inwardly rectifying potassium (GIRK) channels rule synaptic plasticity that supports hippocampal-dependent memory, here we assessed their unknown role in hippocampal oscillatory activity in relation to synaptic plasticity induction.

In alert male mice, pharmacological GIRK modulation did not alter neural oscillations before long-term potentiation (LTP) induction. However, after an LTP generating protocol, both *gain*- and *loss-of* basal GIRK activity transformed LTP into long-term depression, but only specific suppression of constitutive GIRK activity caused a disruption of network synchronization (*δ, α, γ* bands), even leading to long-lasting ripples and fast ripples pathological oscillations.

Together, our data showed that constitutive GIRK activity plays a key role in the tuning mechanism of hippocampal oscillatory activity during long-term synaptic plasticity processes that underlies hippocampal-dependent cognitive functions.

## 1. Introduction

Neuronal oscillations are rhythmic fluctuations of the postsynaptic potentials of a neuronal group (local field potentials, LFP) or a cortical region (electroencephalography, EEG; magnetoencephalography, MEG) that are considered a basic mechanism of neural communication, therefore playing an important role in cognitive and motor processes (Buzsáki et al., 2012; Mysin and Shubina, 2022). Hippocampal-dependent memory formation requires a precise neural activity orchestrated by different brain oscillatory activities, classified according to their frequency (Kalweit et al., 2017). *Delta* oscillations (*δ*, 1-4 Hz) in the CA1 hippocampal area are related to states of non-REM sleep and sensory selection (Girardeau and Lopes-Dos-Santos, 2021; McCormick and Bal, 1997). *Theta* oscillations (*⦵*, 4-8 Hz) arise from different areas such as CA3, CA1 or the medial septum (Colom, 2006; Dragoi et al., 1999) and are associated with cell temporal firing and synaptic plasticity, which are crucial for the encoding and consolidation of spatial memory (Girardeau and Lopes-Dos-Santos, 2021; Hyman et al., 2003). *Alpha* oscillations (*α*, 8-12 Hz) are relevant during working memory formation due to its relationship with attention (Kienitz et al., 2022), while *beta* oscillations (*β*, 12-30 Hz) are related to motor activity (Barone and Rossiter, 2021). *Gamma* oscillations (*ɣ*, 30-100 Hz) play a role in context-association during memory codification and retrieval, crucial for spatial memory (Tort et al., 2009). Finally, *ripples* (100-250 Hz) and *fast ripples* (250-500 Hz) are high frequency oscillations originated in the hippocampus and are involved in memory consolidation (Girardeau and Zugaro, 2011). *Ripples* can be part of physiological and pathological events, while *fast ripples* appear exclusively in pathological cases of altered excitability and excitation/inhibition (E/I) imbalance (Gulyás and Freund, 2015). In addition, oscillations at different frequency bands can interact with each other, which is known as cross-frequency coupling (Salimpour and Anderson, 2019). Among these, *Theta*-*Gamma* Coupling (TGC) is considered a neuronal correlate of learning processes, specifically for spatial learning and the processing of episodic memories (Brooks et al., 2020; Tamura et al., 2017). Furthermore, *theta* and *gamma* oscillations typically occur in an inversely proportional pattern, such that high *theta* power is accompanied by low *gamma* power and *vice versa* (Kalweit et al., 2017).

Furthermore, regarding neural network activity and synaptic plasticity, *theta* oscillations are known as the “*online*” state of the hippocampus since they are necessary for the proper induction of long-term potentiation (LTP) and therefore memory formation (Buzsáki, 2002; Colom, 2006; Nuñez and Buño, 2021). Indeed, it has been suggested that a precise pattern of hippocampal oscillations would be behind the correct generation of the long-term plasticity processes underlying learning and memory capabilities, while at the same time LTP would be a trigger for reorganization of neural network activity (Bikbaev and Manahan-Vaughan, 2007, 2008; Buzsáki and Moser, 2013; Hyman et al., 2003). Thus, neural oscillations and synaptic plasticity are strongly interlinked in the hippocampus.

On the other hand, G-protein-gated inwardly rectifying potassium (GIRK) channels, a family of K^+^ channels that can be activated by different G-protein-coupled receptors (Jeremic et al., 2021b; Lüscher and Slesinger, 2010), have been shown to mediate neuronal synchronization in the firing of pyramidal cells in the hippocampus (Anderson et al., 2021; Dascal, 1997), pointing them out as potential targets for regulating network oscillatory activity (Huang et al., 2005; Kamarajan et al., 2017; Mattis et al., 2014; Sun et al., 2002). The four existing subunits (GIRK1-GIRK4) can array themselves in different combinations to form functional tetrameric channels. In the dorsal hippocampus, GIRK channel conductance is constitutively active (Kim and Johnston, 2015), hyperpolarizing neurons and thus controlling neural excitability (Malik and Johnston, 2017), as well as gating synaptic plasticity processes that support learning and memory formation (Djebari et al., 2021; Malik and Johnston, 2017). Furthermore, GIRK activity has been also linked to alterations in *theta* (Kamarajan et al., 2017) and *gamma* (Johnston et al., 2014; Pietersen et al., 2009) oscillations during reward processing, and might be part of the neurogenetic basis of cognitive dysfunctions in addiction and other disorders (Kamarajan et al., 2017). Even *ripples* are regulated by GIRK activity in the dorsal hippocampus, as its blockage increased *sharp wave-ripples* (*SPW-Rs*)(Trompoukis et al., 2020). However, the role of GIRK activity in hippocampal oscillations during long-term synaptic plasticity processes remains largely unknown.

Thus, there is a strong and finely tuned relationship between synaptic plasticity and oscillatory activity in the hippocampus for proper memory acquisition. GIRK channels have a crucial ruling function in synaptic plasticity underlying memory formation and retrieval (Djebari et al., 2021). Therefore, the aim of the present study was to determine, their role in dorsal hippocampal oscillatory activity before and after the induction of long-term synaptic plasticity in alert animals to further understand the mechanisms by which GIRK channels may modulate hippocampal function.

## 2. Materials and methods

### 2.1. Animals

Male C57BL/6 adult mice (*n* = 57; 3-5-month-old; 28-35 g) were used (Charles River). Before surgery, animals were housed in groups of 5 per cage, whereas after surgery animals were housed individually. They were kept on 12 h light/dark cycles with constant temperature (21 ± 1 ºC) and humidity (50 ± 7%). Food and water were available *ad libitum*.

All experimental procedures were reviewed and approved by the Ethical Committee for Use of Laboratory Animals of the University of Castilla-La Mancha (PR-2015-01-01 and PR-2018-05-11) and conducted according to the European Union guidelines (2010/63/EU) and the Spanish regulations for the use of laboratory animals in chronic experiments (RD 53/2013 on the care of experimental animals: BOE 08/02/2013).

### 2.2. Drugs

All chemicals used in this study were purchased from Abcam (Cambridge, UK) and dissolved in phosphate-buffered saline (PBS). To pharmacologically modulate GIRK channels, a blocker and a selective activator were used. Tertiapin Q (TQ; 1.2 μg/μL) blocks GIRK channels containing subunits 1 and 4, while ML297 (0.5 μg/μL) selectively opens GIRK1-containing channels. Concentrations were determined by preliminary tests and based on previous reports (Djebari et al., 2021; Kotecki et al., 2015). Animals were randomly assigned to experimental groups, and they received a 3 μL intracerebroventricular *(icv*.) injection of either TQ, ML297 or vehicle (PBS) through an injection cannula inserted in a chronically implanted guide cannula (see details below) using a Hamilton syringe and a pump at a rate of 0.5 μL/min. *Icv*. injections were performed in alert freely moving mice as described elsewhere (Djebari et al., 2021; Sánchez-Rodríguez et al., 2020).

### 2.3. Surgery

For chronic recording and drug injection, mice underwent surgery to be implanted with recording and stimulating electrodes and a guide cannula (Figure 1A). To do so, mice were anesthetized with 4–1.5% isoflurane (induction and maintenance, respectively; ISOFLO, Proyma S.L.) delivered using a calibrated R580S vaporizer (RWD Life Science; flow rate: 0.5 l/min O2) as previously described (Djebari et al., 2021).

**Figure 1.**
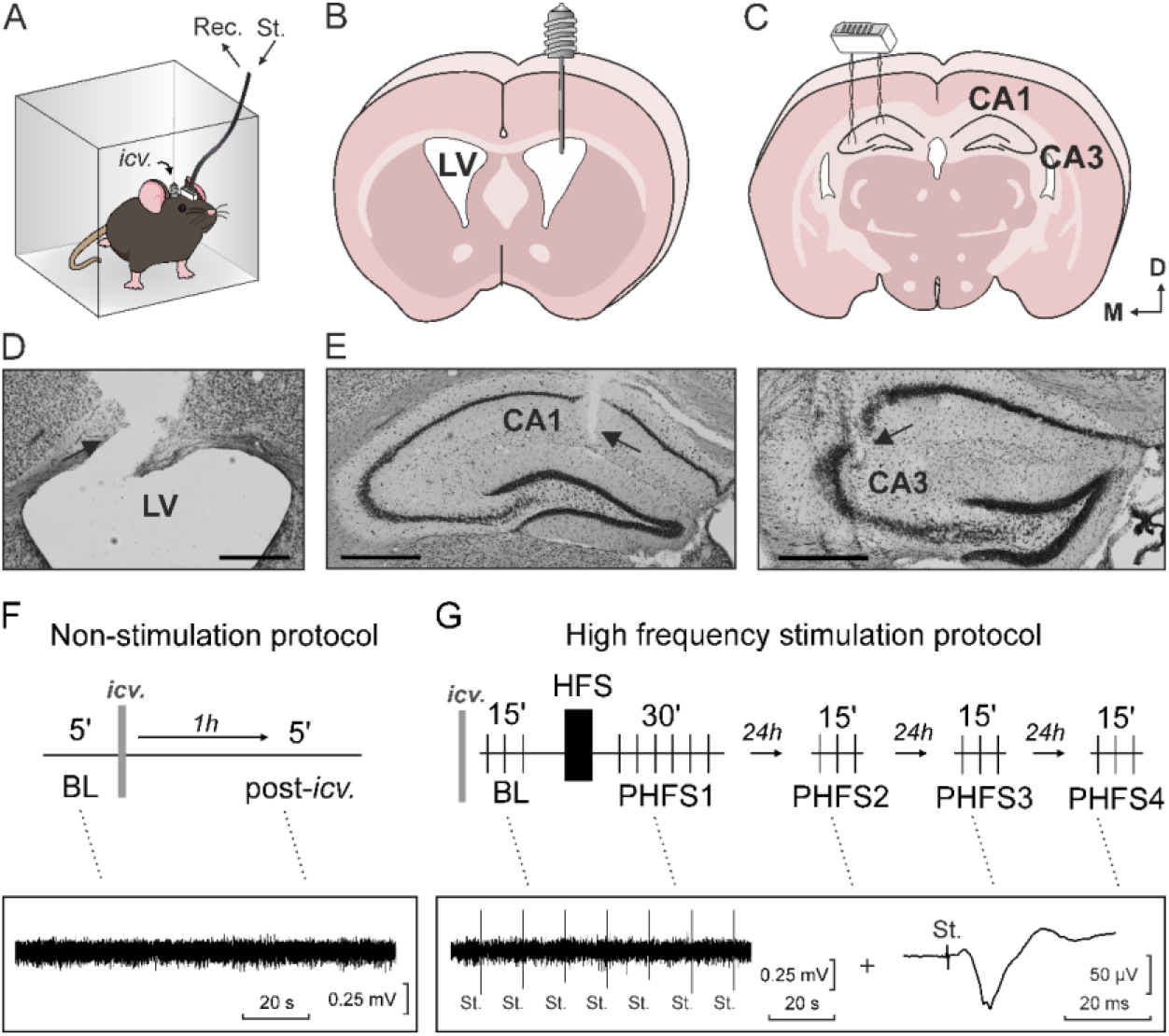
Experimental design. **(A)** For all the experiments, mice were placed in a 9.5 × 9.5 × 7 cm small box where they could freely move while intracerebroventricular (*icv*.) injections, recording and/or stimulation were performed. **(B)** Location of the stainless-steel guide cannula implanted for *icv*. drug administration, on the left ventricle, contralaterally to recording/stimulation electrodes, in order to preserve the functionality of CA3–CA1 synapse. **(C)** Location of electrodes surgically implanted in CA1 (recording) and CA3 (stimulating) for chronic fEPSPs and LFPs recordings. **(D)** Histologic verification of cannula position (black arrow). Scale bar: 500 µm. **(E)** Histologic verification of CA1 (left) and CA3 (right) electrodes position (black arrows). Scale bars: 500 µm. **(F)** Schematic diagram of the non-stimulation protocol, in which LFPs were recorded pre- (baseline, BL) and post-*icv*. treatment without applying any external stimulation. **(G)** Schematic diagram of the high frequency stimulation protocol (HFS) used, in which both fEPSPs (with single electrical pulses applied every 15 sec) and LFPs were recorded pre- and post-HFS stimulation. LV, lateral ventricle; Rec. recording; St, stimulation; D, dorsal; M, medial.

Two bipolar electrodes, made from 50 μm Teflon-coated tungsten wire (Advent Research Materials, UK), were implanted in each animal at the right hippocampus (Figure 1B). The stimulating electrode was aimed at the Schaffer collateral-commissural pathway in the CA3 area (2 mm lateral and 1.5 mm posterior to bregma; depth from brain surface, 1.0–1.5 mm). The recording electrode was aimed at the *stratum radiatum* underneath the CA1 area (1.2 mm lateral and 2.2 mm posterior to bregma; depth from brain surface, 1.0–1.5 mm) (Paxinos and Franklin, 2004). The final position of the hippocampal electrodes was verified by Nissl staining after experimental completion (Figure 1E). A silver wire (0.1 mm) was affixed to the skull as ground. Both electrodes and ground were weld to a 6-pin socket (Mouser Electronics, US), which was attached to the skull using dental cement.

For *icv*. administration of the studied drugs, animals were also implanted with a blunted, stainless steel, 26-G guide cannula (Plastics One, US) in the left ventricle (Figure 1B; 1 mm lateral and 0.5 mm posterior to bregma; depth from brain surface, 1.8 mm) (Paxinos and Franklin, 2004). The cannula was implanted contralaterally to the bipolar electrodes to preserve the physiological properties of the CA3-CA1 synapse in the right hippocampus. Diffusion of *icv*. injections to both dorsal hippocampi has been shown elsewhere (Djebari et al., 2021; Sánchez-Rodríguez et al., 2020). The final position of the cannula was also confirmed by Nissl staining (Figure 1D). Mice were allowed at least a week for recovery before experimental procedures.

### 2.4. *In vivo* electrophysiological recordings

LFP activity and field excitatory postsynaptic potentials (fEPSPs) were recorded from the hippocampal CA1 area of alert behaving mice with Grass P511 differential amplifiers through a high-impedance probe (2 × 1012 Ω, 10 pF).

Firstly, to evaluate GIRK modulation effect on hippocampal oscillatory activity, the animal was placed in a small box (9.5 × 9.5 × 7 cm; Figure 1A) and LFPs were recorded, without applying any type of stimulation, during 5 min before (pre-treatment values) and 1 h after *icv*. injections (Figure 1F), from which up to 3 min of recordings, free of unwanted artifacts, were selected for subsequent spectral analysis using the Spike2 software.

Secondly, to determine GIRK modulation effect on network function after high-frequency stimulation (HFS)-induced synaptic plasticity, both fEPSPs and LFPs were collected during 15 min prior to LTP induction as a baseline (BL), stimulating with single 100 μs, square, biphasic pulses elicited at 0.05 Hz, as obtained previously (Djebari et al., 2021). The stimulus intensity was selected according to two criteria: 1) ∼35% of its asymptotic value, and 2) a second stimulus, presented 40 ms after, evoked a larger (≥150%) synaptic field potential than the first one (Bliss and Gardner-Medwin, 1973). After BL, a HFS protocol was applied, consisting of five 100 Hz, 100 ms trains of pulses at a rate of 1/s repeated 6 times, at intervals of 1 min, reaching a total of 300 pulses in each HFS session (Sánchez-Rodríguez et al., 2017). To avoid evoking large population spikes and/or the appearance of cortical seizures, the same stimulus intensity was used during BL, HFS and post-HFS (PHFS) sessions. After HFS protocol application for LTP induction, fEPSPs and LFPs were recorded during 30 min that same day (PHSF-1) and during 15 min on three consecutive days (PHFS-2, -3 and -4; Figure 1G). All fEPSP amplitude data obtained from the CA1 hippocampal region during the HFS session and afterwards were normalized using BL fEPSP values collected on the first day as 100%. The power spectrum of the hippocampal activity was computed with the Spike2 software, using the average of all 15 sec windows between pulses, starting 1 sec after the stimulus (Figure 4A).

Frequency bands analyzed in both sets of experiments were: *delta* (*δ*, 1-4 Hz), *theta* (*θ*, 4-8 Hz), *alpha* (*α*, 8-12 Hz), *beta* (*β*, 12-30 Hz), *gamma* (*γ*, 30-100 Hz), *ripples* (100-250 Hz) and *fast ripples* (250-500 Hz) (Kalweit et al., 2015).

### 2.5. Data collection

Recordings were stored digitally on a computer through an analog/digital converter (CED 1401 Plus). Data were analyzed off-line for quantification of fEPSP amplitude and LFPs, using the Spike2 (CED) program. Spectral analysis is one of the standard methods used for quantification of LFP and reflects the distribution of signal power over different frequencies. To do so, Fourier Transform (FT) was used, a mathematic algorithm that transform the signal into frequency-dependent functions. Thus, the power spectrum provides data from each frequency band, allowing to study different bands and their relationships with different physiological processes.

### 2.6. Statistical analysis

Data were represented as the mean ± SEM and analyzed by one-way or two-way ANOVA, with time and treatment as within- and between-subjects factors, respectively. For repeated measures two-way ANOVA, Greenhouse-Geiser correction was applied and indicated in the text when sphericity was not assumed. Specific information for each experiment can be found in the Results section. Statistical significance was set at *p* < 0.05. Statistical analysis was performed using GraphPad Prism software (version 8.3.1). Final figures were prepared using CorelDrawX8 Software.

## 3. Results

### 3.1. GIRK modulation does not affect hippocampal network activity in basal conditions

GIRK channels dysfunction has been related to different central nervous system pathologies and cognitive deficits caused by an imbalance in neuronal excitability (Jeremic et al., 2021b), such as epilepsy (Huang et al., 2018; Mazarati et al., 2006), Down syndrome (Cooper et al., 2012; Reeves et al., 1995; Sago et al., 1998) or Alzheimer’s disease, which can be associated with multiple alterations in oscillatory network activity (Mayordomo-Cava et al., 2020; Mayordomo-Cava et al., 2015; Nava-Mesa et al., 2013; Sánchez-Rodríguez et al., 2017). Therefore, we decided to study the role of these channels in the oscillatory activity of the dorsal hippocampal CA1 region by analyzing the LFPs recordings before and after *icv*. administration of GIRK channel modulators: either the channel blocker TQ (*n* = 11), or the selective activator of the channel ML297 (*n* = 9), or PBS as control vehicle (*n* = 15) (Figure1F; Figure 2). To do so, the following frequency bands were analyzed: *δ* (1-4 Hz), *⦵* (4-8 Hz), *α* (8-12 Hz), *β* (12-30 Hz), *γ* (30-100 Hz), *ripples* (100-250 Hz) and *fast ripples* (250-500 Hz). Finally, their power during the post-injection session were as percentage respecting the value found during the pre-injection session.

**Figure 2.**
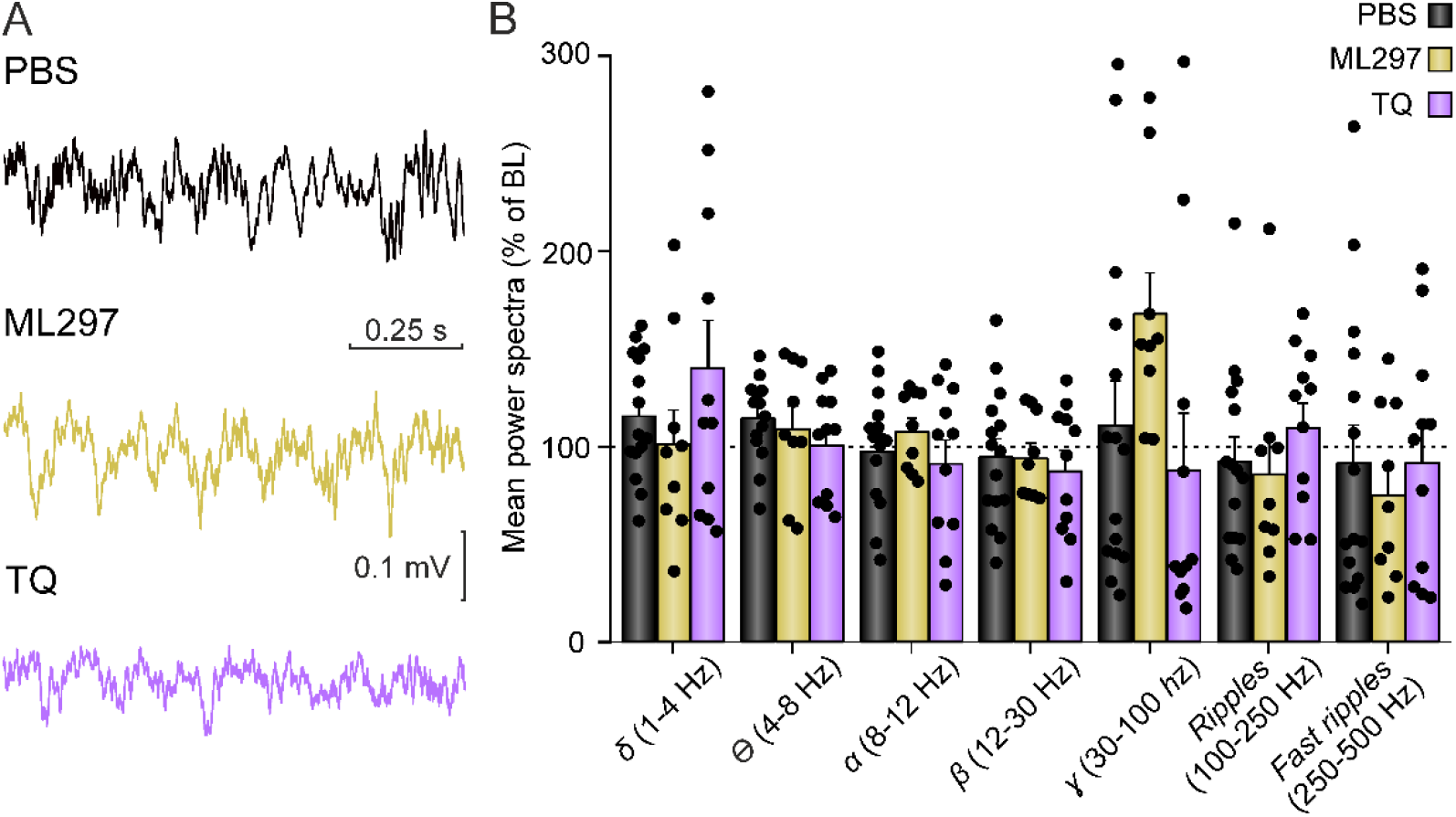
Pharmacological modulation of GIRK function does not affect hippocampal oscillatory activity in non-stimulating conditions. **(A)** Representative examples of LFPs recorded from the CA1 region of the dorsal hippocampus in alert mice after treatment with GIRK modulators ML297 (channel opener) and TQ (channel blocker). **(B)** Histograms of the spectral power of LFP activities recorded in the CA1 region after PBS (control vehicle), ML297 or TQ injections. The dashed line represents the percentage of baseline (BL, 100%) recordings (pre-treatment). Note that pharmacological modulation of GIRK channel did not alter the values of selected spectral bands for neither *δ, ⦵, α, β, ɣ, ripples* nor *fast ripples* oscillations. Data is expressed as mean ± SEM and as a percentage of baseline recordings (BL, 100%).

Results showed that there were no significant differences after *icv*. injection of neither the opener nor the blocker of GIRK channels in any of the frequency bands analyzed (Figure 2A-B): *δ* (F_(2,32)_ = 1.201, *p* = 0.345), *θ* (F_(2,32)_ = 0.847, *p* = 0.438), *α* (F_(2,32)_ = 0.664, *p* = 0.522), *β* (F_(2,32)_ = 0.187, *p* = 0.831), *γ* (F_(2,32)_ = 2.303, *p* = 0.116), *ripples* (F_(2,32)_ = 0.722, *p* = 0.493) and *fast ripples* (F_(2,32)_ = 0.235, *p* = 0.792). This result suggests that pharmacological modulation of GIRK channels is not able to modify CA1 hippocampal oscillatory activity under basal conditions, i.e. in the absence of any artificial stimulation.

### 3.2. Constitutive GIRK activity is needed for proper hippocampal synaptic plasticity

However, as aforementioned, LTP generation in the hippocampus is accompanied by changes in neuronal oscillations (Bikbaev and Manahan-Vaughan, 2007). Given that previous work from our group showed that GIRK channel pharmacological modulation inhibits LTP both *in vivo* and *in vitro* (Djebari et al., 2021; Sánchez-Rodríguez et al., 2017), we aimed to study the role of GIRK channels in the oscillatory activity of the dorsal hippocampus after a LTP induction protocol (see methods section for details). To do so, first, we studied the effect of constitutive GIRK channel activity modulation on LTP induction by applying HFS at the Schaffer collateral pathway to induce LTP in the CA1 region; and compared the evolution of fEPSPs in the different groups and at different timepoints (Figure 1G). Briefly, fEPSPs were obtained by single pulse stimulation at 0.05Hz (i.e, every 15 sec) at pre- (baseline, BL) and at four post-HFS periods (following HFS on day 1, PHFS-1; and 24, 48 and 72h post-HFS on days 2, 3 or 4 respectively (PHFS-2, -3 or -4) (Figure 3).

**Figure 3.**
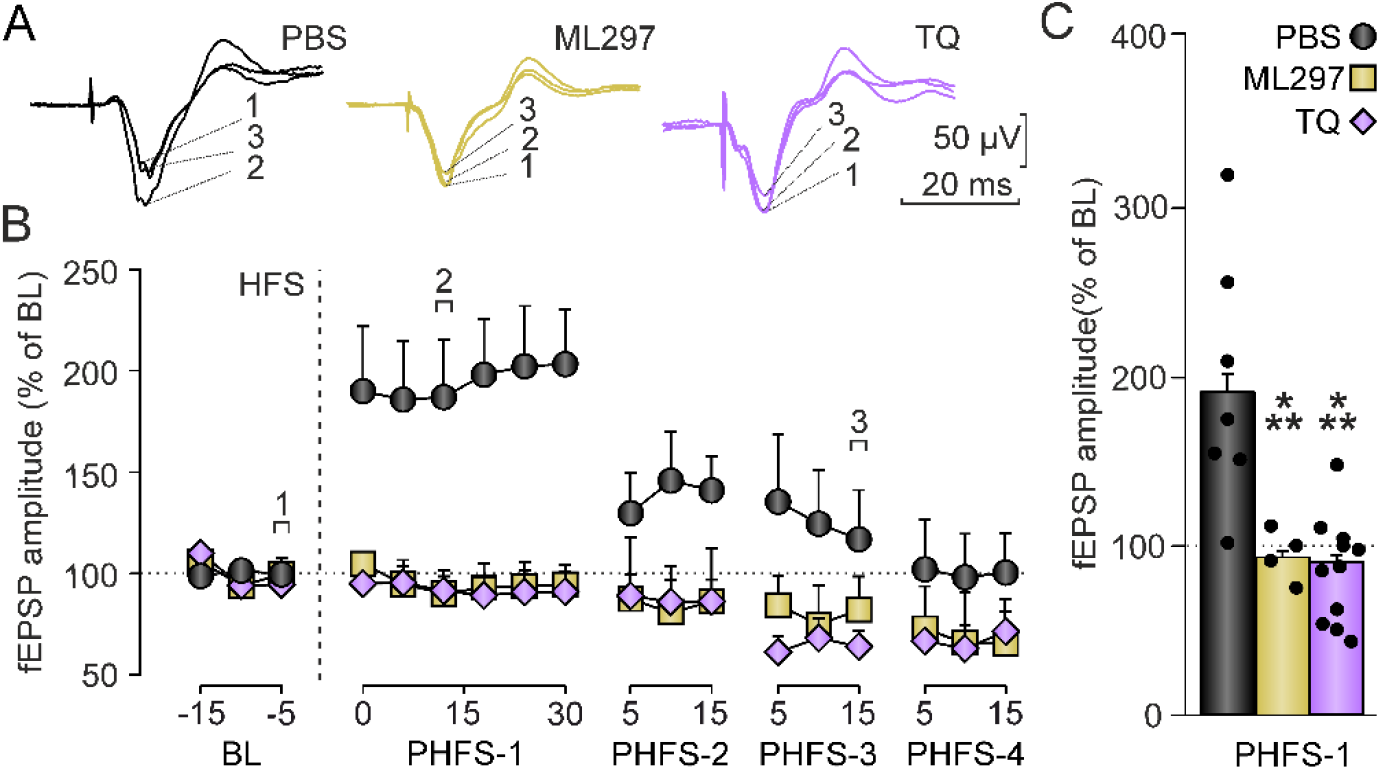
Constitutive GIRK activity is needed for proper hippocampal synaptic plasticity *in vivo*. **(A)** Representative examples of fEPSPs (averaged 5 times) evoked at the CA3–CA1 synapse by single stimulation collected after *icv*. drug administration but prior to HFS (1, BL), following HFS (2) and 48 h after HFS (3) in mice injected with vehicle (PBS, control), GIRK channel opener ML297 or channel blocker TQ. **(B)** Evolution of fEPSPs recorded in alert mice of each experimental group during 15 min prior to HFS (BL), during 30 min after LTP induction that same day (PHFS-1) and during 15 min on three consecutive days (PHFS-2, -3 and -4). Data is expressed as the mean amplitude ± SEM and as a percentage of BL recordings (100%) **(C)** Bars illustrate fEPSPs amplitude to show potentiation level (mean ± SEM) after PHFS-1. *** *p* < 0.001 *vs*. PBS.

Here, our results showed a significant treatment effect (Figure 3C; F_(2,19)_ = 10.956, *p* > 0.01) between controls and both treated groups. The stimulation protocol induced a robust LTP in CA3-CA1 synapses of vehicle-injected mice (Figure 3A-C; 194 ± 11% of BL; *n* = 7) during the 30 min following HFS (F_(8,48)_ = 9.061, *p* < 0.001). However, LTP induction was not achieved after pharmacological GIRK channel activity increase nor reduction through *icv*. injections of ML297 (F_(8,48)_ = 0.571, *p* = 0.791; *n* = 4) or TQ (F_(1.73,17.27)_ = 1.184, *p* = 0.323, Greenhouse-Geiser correction; *n* = 11), respectively (Figure 3A-C). Moreover, 72 h after HFS, while the amplitude of CA1 fEPSPs recovered BL values in control animals (100 ± 11% of BL), both modulations induced a noticeable depression of the synaptic response (Figure 3B; ML297: 68 ± 12% of BL; TQ: 68 ± 5% of BL), thus showing an HFS-induced long-term depression (LTD).

### 3.3. Constitutive GIRK activity supports network function after HFS-induced synaptic plasticity

Finally, to examine the possible changes in hippocampal oscillations that might accompany the alterations observed in synaptic plasticity, LFPs were also recorded during BL and the same post-HFS periods described above (PHSF-1, -2, -3 and -4) in all the 15 sec windows between single pulses (Figure 1G and Figure 4A).

**Figure 4.**
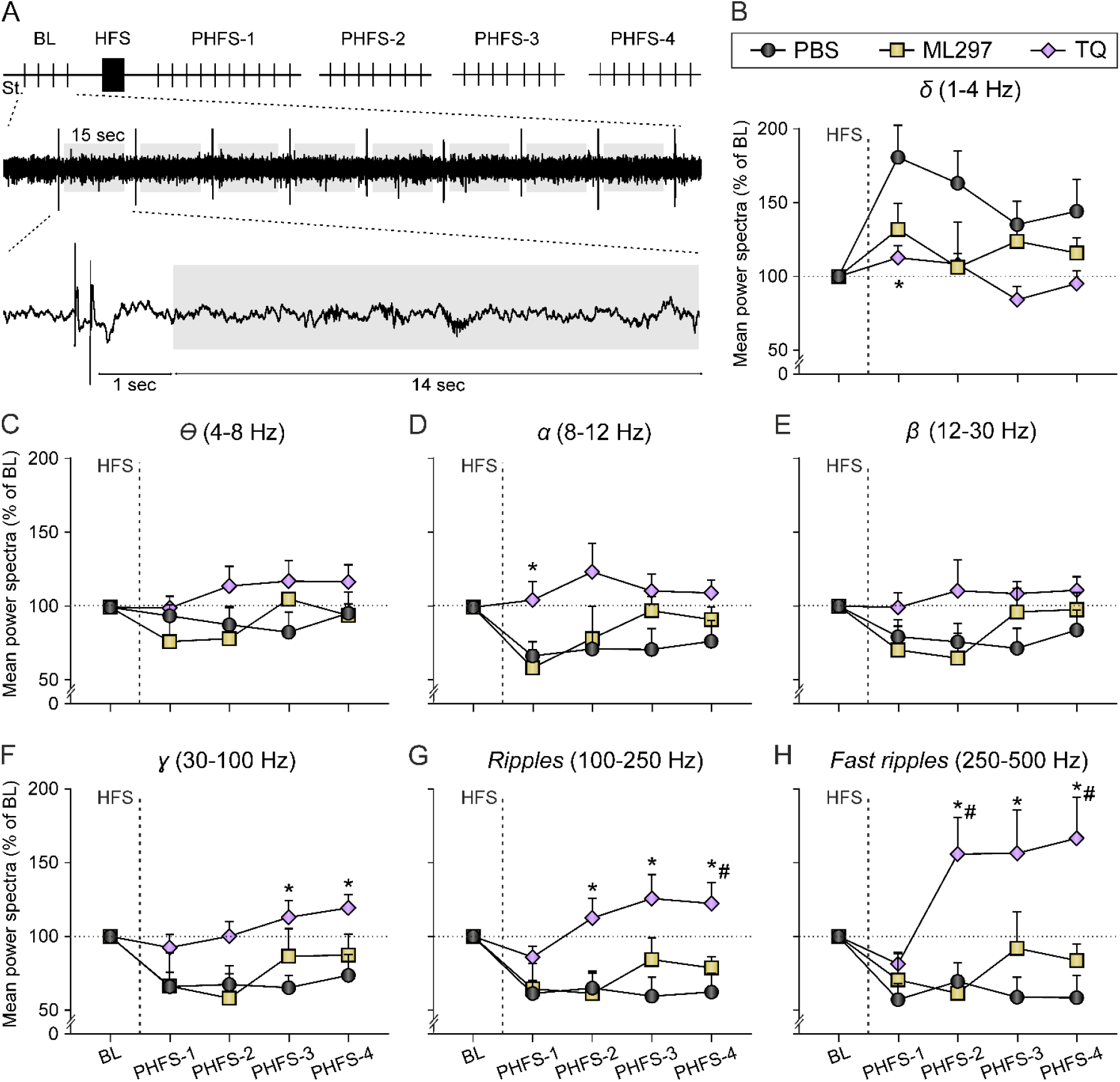
Constitutive GIRK activity supports network function after HFS-induced synaptic plasticity. **(A)** Representation of the protocol used to quantify LFPs in all the 15 sec windows between pulses during BL and the different post-HFS sessions (PHFS-1, -2, -3 and -4). Average of spectral power of frequency intervals corresponding to *delta* **(B)**, *theta* **(C)**, *alpha* **(D)**, *beta* **(E)**, *gamma* **(F)**, *ripples* **(G)** and *fast ripples* **(H)** bands. Data is expressed as mean ± SEM and as a percentage of baseline recordings (BL, 100%). * *p* < 0.05 *vs*. PBS (vehicle); # *p* < 0.05 *vs*. ML297.

Regarding low frequency bands, data showed a significant treatment effect in the *δ* band (Figure 4B; F_(2,20)_ = 8.649, *p* = 0.002). *Post-hoc* analysis revealed differences between mice injected with TQ and controls during PHFS-1. Thus, *δ* frequency was increased in control animals right after LTP induction (during PHFS-1) but not in TQ treated mice, suggesting a role of GIRK channels in *δ* rhythm generation during HFS. Furthermore, there was a significant time effect (F_(2.89,56.38)_ = 4.473, *p* = 0.0075, Greenhouse-Geiser correction), since the initial increment progressively declined during the following days (PHFS-2, -3 and -4).

On the other hand, neither increasing nor decreasing GIRK-dependent signaling in the dorsal hippocampus significantly altered the *⦵* band after a HFS protocol applied to induce LTP (Figure 4C; F_(2,19)_ = 2.032, *p* = 0.1586). The analysis of the *α* band showed a significant treatment effect (Figure 4D; F_(2,19)_ = 7.140, *p* = 0.0049), specifically during PHFS-1, in which *alpha* frequency was transiently decreased in control and ML297-injected mice but stayed approximately at BL levels in TQ-injected mice. Moreover, the *β* band was also altered due to treatment (Figure 4E; F_(2,19)_ = 6.665, *p* = 0.0064). However, *post-hoc* analysis did not show differences in any specific session. Data also showed a significant treatment effect in the *ɣ* band (Figure 4F; F_(2,19)_ = 9.764, *p* = 0.0012). The effect was observed during PHFS-3 and PHFS-4, between control animals and TQ treated mice.

*Gamma* frequency was decreased in control mice after LTP induction, yet in the TQ group was progressively enhanced, reaching its peak at PHFS-4. As it happened with the *delta* band, a significant time effect was also detected (F_(3.10,55.84)_ = 2.951, *p* = 0.0388, Greenhouse-Geiser correction).

Regarding the high frequency bands, a significant treatment effect was observed on the *ripples* band (Figure 4G; F_(2,19)_ = 10.411, *p* = 0.0009). *Post-hoc* analysis revealed differences between TQ animals and controls, as both controls and ML297 animals showed a reduction of this frequency after LTP induction, while TQ mice displayed an increment, specifically during PHFS-2, -3 and -4. Finally, when analyzing the *fast ripples* band a significant treatment effect was detected (Figure 4H; F_(2,19)_ = 8.008, *p* = 0.003). Once again, TQ-treated mice presented an increase in this band during PHFS-2, -3 and -4, that was not observed in control or ML297 mice.

Hence, all these alterations in the power spectra induced by the lack of GIRK activity after an HFS protocol, but not under non-stimulation conditions, suggest a differential role for GIRK channels in neural oscillations and long-term synaptic plasticity processes that underlie learning and memory.

## 4. Discussion

During hippocampal-dependent memory processing, and the underlying long-term synaptic plasticity generation, significant changes in neuronal oscillation activity occur (Abubaker et al., 2021; Bikbaev and Manahan-Vaughan, 2008, 2017), and our results show that GIRK channel activity have a relevant role in these interlinked processes.

### 4.1. GIRK activity modulation does not alter basal network oscillations at dorsal hippocampal CA1 region

Given the link between oscillatory activity, LTP and GIRK channels, we started by pharmacological modulation of GIRK channel function in dorsal hippocampus of alert mice without applying any artificial external stimulation, to study their contribution to oscillatory activity. Our results showed no significant changes in any of the frequency bands considered in this study. Nevertheless, other authors have found different results. For instance, Tatard-Leitman et al. (2015) showed an increase in *⦵* (4-12 Hz), *β* (13-30 Hz) and *ɣ* (30-80 Hz) bands in a NMDA-R1 knocked-out mice model that exhibited a decreased expression of GIRK2 subunit in the synaptic membrane. However, since several hippocampal synaptic pathways are affected by this manipulation, it cannot be concluded that these changes are exclusively due to the GIRK channel down-regulation. Also, the pharmacological blockade of GIRK channels by TQ increased the rate of occurrence of *SPW-Rs* (110-200 Hz) in dorsal hippocampal slices (Trompoukis et al., 2020). Similarly, GIRK2 constitutive KOs show increased susceptibility to seizures (Blednov et al., 2001; Pravetoni and Wickman, 2008; Victoria et al., 2016), which could be related to this enhancement in the amplitude of neural oscillations. In contrast, in our approach, *icv*. application of a GIRK blocker did not elicit any signs of epileptiform activity in the hippocampus (Djebari et al., 2021). Furthermore, previous work from our laboratory showed that, while ML297 injection had no effect on *⦵* and *ɣ* spectral bands, this GIRK opener was able to rescue control values when it was applied along with Amyloid-*β* peptide (a toxic strongly related to increased excitability neuropathology in Alzheimer’s disease (Jeremic et al., 2021a)), since both *⦵* and *ɣ* are known to be dramatically altered by hyperexcitability in Alzheimer’s disease animal models (Sánchez-Rodríguez et al., 2017). Thus, the impact of GIRK modulation on dorsal hippocampal oscillatory activity *in vivo* seems to be dependent on hippocampal excitability levels, being of greater importance in pathological conditions involving an Excitatory/Inhibitory (E/I) imbalance.

### 4.2. Constitutive GIRK activity supports synaptic plasticity and network function after HFS protocol

It has been reported that mice lacking GIRK2 in forebrain pyramidal neurons (CaMKII-Cre(+):GIRK2^flox/flox^) exhibited a marked depletion of LTP as well as an enhanced LTD and blunted LTP depotentiation (Victoria et al., 2016). Additionally, GIRK channels are critical for LTP of synaptic inhibition, a form of synaptic plasticity that prevents saturation of synaptic potentiation and increases the flexibility and storage capacity of neuronal circuits (Huang et al., 2005; Sánchez-Rodríguez et al., 2019). In addition, both GIRK opening and blockade disrupted LTP induction. In the case of ML297, an intense pyramidal neurons hyperpolarization and subsequent insufficient activation of NMDA receptors (NMDARs) might explain LTP impairment (Djebari et al., 2021), while TQ would increase GABA release producing the inhibition of pyramidal neurons (Davies et al., 1991). Both mechanisms would explain the results observed in the dorsal hippocampus *in vivo*, showing a critical role for GIRK channel conductance in regulating LTP/LTD induction threshold and subsequent memory processes (Djebari et al., 2021). Moreover, HFS-induced LTD could be explained, at least partially, by the Bienenstock, Cooper and Munro (BCM) theory, which states that changes in neuronal excitability, with GIRK channels being of great importance in its control in the hippocampus, and thus in hippocampal activity levels, can alter the induction threshold of long-term plasticity processes (Keck et al., 2017). However, another possible explanation could be found when neuronal oscillations that occur during synaptic plasticity generation are explored, as we did here.

Consequently, the next step was to analyze whether pharmacological GIRK channel activity modulation could influence hippocampal oscillations that pair with synaptic plasticity events. Our results indicated that, while GIRK activity enhancement by ML297 administration did not significantly modify any of the frequency bands analyzed, the reduction of GIRK function using TQ caused imbalances in almost all neuronal oscillations during synaptic plasticity induction and maintenance. In dorsal CA1, GIRK channels are constitutively active and cause a decrease in excitability, contributing to the E/I balance (Kim and Johnston, 2015). Increasing GIRK activity above physiological basal values with ML297 would increase the threshold for LTP induction causing its disruption (Cummings et al., 1996; Djebari et al., 2021). According to our present results, this mechanism would be independent of hippocampal oscillations. On the other hand, decreasing GIRK constitutive activity with TQ, and therefore eliminating some of the necessary basal inhibitory transmission required for proper hippocampal function, seems to alter the oscillatory activity needed for proper LTP induction (Bikbaev and Manahan-Vaughan, 2008). Thus, both function gain and loss of GIRK activity impair LTP, through two different pathways.

To deepen the understanding of the involvement of GIRK channel conductance in hippocampal network activity following LTP induction, we will first focus on cross-frequency coupling, a fundamental feature of oscillatory activity that is a critical component of physiological cognitive function (Canolty and Knight, 2010; Lisman and Jensen, 2013). TGC is the most studied coupling, and several works have demonstrated its involvement in memory formation processes (Canolty and Knight, 2010; Lisman and Jensen, 2013), suggesting that *theta* and *gamma* oscillations could act as an intrinsic mechanism for LTP to occur (Bikbaev and Manahan-Vaughan, 2008). In this regard, Bikbaev and Manahan-Vaughan (2008) showed that LTP is associated with a very specific pattern of TGC changes: an increase in *theta* power and a transient increase (up to 100 ms after high-frequency tetanisation) followed by a decrease in *gamma* power. Although our experimental procedure, in which we averaged the spectral power over the entire session (30 or 15 min), did not allow us to observe such short transient variations in the different frequency bands, our results indicate a decrease in *ɣ* power after LTP induction in control and ML297-treated animals that lasts at least 4 days post-HFS. Interestingly, the blockade of GIRK channels by TQ progressively increases this frequency, being significantly higher on the last 2 days and coinciding with the depression of the fEPSP amplitude observed in these mice, which may be partially underlying the HFS-induced LTD described here and elsewhere (Djebari et al., 2021). In this line, a link between GIRK channels decreased expression and an increased baseline hippocampal *ɣ* power (Tatard-Leitman et al., 2015) has been previously proposed. Surprisingly, we did not observe any changes in the amplitude of *⦵* power, neither in control mice nor after GIRK modulation. These differences with Bikbaev and Manahan-Vaughan (2007) results may be due to the stimulating protocol to induce LTP since we used a HFS protocol (Gruart et al., 2006), while they applied a *theta*-burst stimulation (TBS) protocol. These two protocols are known to trigger distinct intracellular signaling cascades, which elicit different outcomes for LTP (Hegemann and Abraham, 2021; Liu et al., 2020; Mastrolia et al., 2021), hence they may be affecting the *⦵* power differently as well.

Less is known about how other rhythms such as *delta, alpha* or *beta* are modulated after a plasticity mechanism is triggered. Our results showed an enhancement of *δ* power as well as a reduction of *α* power after *in vivo* HFS in control animals, that were reversed in TQ-treated mice. Similarly, other work had described this suppression of the *δ* frequency when LTP is hindered in a rat model of psychosis (Kalweit et al., 2017). Furthermore, *delta*-*alpha* coupling influence TGC and serve as a gating mechanism to prioritize relevant information in a working memory task (Leszczyński et al., 2015). Thus, the alterations on these two frequencies may be related to the learning and memory deficits previously reported when GIRK activity decreases (Djebari et al., 2021; Mett et al., 2021). Regarding *β* oscillations, our results did not show any specific differences, which could be due to the relationship of this frequency with motor activity (Barone and Rossiter, 2021), since in our animal model were restrained in a small box and not performing any behavioral task.

*SPW-Rs*, which are modulated by 5-HT_1A_ in the hippocampus, have also been related to memory consolidation and can be triggered by the same HFS protocols that induce LTP (ul Haq et al., 2016). Given the described relationship between GIRK channels and 5-HT_1A_ receptors (Jeremic et al., 2021b) and their role in the modulation of hippocampal *gamma* oscillations (Johnston et al., 2014), the evaluation of *ripples* and *fast ripples* gains special interest. Here, we found that GIRK channels blockade did increase both *ripples* and *fast ripples* power after a HFS protocol, even days after synaptic plasticity induction. It has been already described that GIRK channel blockade by TQ increases the rate of *ripples* occurrence in dorsal hippocampal slices (Trompoukis et al., 2020), as we find here in alert animals. *Ripples* have been shown to be critically involved in the process of memory consolidation during sleep, and they are generated by the disinhibition of pyramidal cells (Evangelista et al., 2020). Thus, we observed low levels of this oscillation frequency in control animals, consistently with the fact that they were awake, and incremented levels in TQ-treated mice, as blocking GIRK activity would lead to hippocampal disinhibition. Furthermore, loss of inhibition turns *ripples* into *fast ripples*, as our results show as well, which are pathological oscillations related to epileptic activity (Gulyás and Freund, 2015), where GIRK channels have been shown to play a critical role (Jeremic et al., 2021b).

## 5. Conclusions

In summary, neuronal oscillations are crucial for supporting normal information processing and storage in the hippocampus, and LTP and oscillatory activity are reciprocally interlinked to contribute to memory formation (Bikbaev and Manahan-Vaughan, 2007). Here, we found that the suppression of GIRK constitutive activity in the dorsal hippocampus causes a disruption of the correct network synchronization, leading to pathological oscillations (*fast ripples)* during a long-term synaptic plasticity event but not under basal conditions (without artificial stimulation). Consequently, all these alterations in spectral power may account for the LTP impairment and memory deficits previously reported (Djebari et al., 2021; Sánchez-Rodríguez et al., 2020; Sánchez-Rodríguez et al., 2019; Sánchez-Rodríguez et al., 2017) and shed light on how GIRK channels contribute to synaptic and network processes that supports hippocampal-dependent cognitive capabilities.

## Acknowledgments

We thank José M. Gonzalez Martín, María Sánchez Enciso, and José A. Santos for their excellent technical assistance.

## Author Contributions

Conceptualization, LJD and JDNL; Investigation, STC; Formal analysis, AM, AC and SD; Resources, JMDG, AG, LJD and JDNL; Writing - original draft, AC and SD; writing – review and editing, AC, SD, LJD and JDNL; Funding acquisition, LJD and JDNL.

## Data statement

The data reported in this paper is available from the corresponding authors upon request.

